# Target Recycling Amplification Process for Digital Detection of Exosomal MicroRNAs Through Photonic Resonator Absorption Microscopy

**DOI:** 10.1101/2022.12.07.519503

**Authors:** Xiaojing Wang, Skye Shepherd, Nantao Li, Congnyu Che, Tingjie Song, Yanyu Xiong, Isabella Rose Palm, Bin Zhao, Manish Kohli, Utkan Demirci, Yi Lu, Brian T. Cunningham

**Author notes:** These authors contributed equally to this work (Xiaojing Wang, Skye Shepherd).

## Abstract

Exosomal microRNAs (miRNAs) have considerable potential as pivotal biomarkers to monitor cancer development, dis-ease progression, treatment effects and prognosis. Here, we report an efficient target recycling amplification process (TRAP) for the digital detection of exosomal miRNAs using photonic resonator absorption microscopy (PRAM). Through toehold-mediated DNA strand displacement reactions, we achieve multiplex digital detection with sub-attomolar sensitivity in 20 minutes, robust selectivity for single nucleotide variants, and a broad dynamic range from 1 aM to 1 pM. We then applied our TRAP system to quantify miRNA in exosomal total RNAs isolated from human cancer cell lines. Compared with traditional qRT-PCR methods, TRAP showed similar accuracy in profiling exosomal miRNAs derived from cancer cells, but also exhibited at least 31-fold and 61-fold enhancement in the limits of miRNA-375 and miRNA-21 detection, respectively. The TRAP approach is ideal for exosomal or circulating miRNA biomarker quantification, where the miRNAs are present in low concentrations or sample volume, with potentials for frequent, low-cost, and minimally invasive point-of-care testing.

## Introdution

Exosomal microRNAs (miRNAs) sequestered within extracellular vesicles play diverse roles in biological processes, including cell-cell communication, cell proliferation, and inflammatory response.^1–2^ miRNAs participate in post-transcriptional regulation of gene expressions. As a result, inappropriate miRNA release from exosomes can result in the development of cardiovascular diseases^3^ and cancers. On this basis, exosomal miRNAs are recognized as pivotal biomarkers for diagnosing cancer development and monitoring the progression of diseases, prognostication and determining therapy outcomes.^4–5^ However, exosomal miRNAs may be present in extremely low concentrations, which is a current barrier to utilizing exosomal miRNAs as biomarkers. For exosomes isolated from cells or plasma, there can be far less than a single miRNA per exosome on average, even for the most abundant target sequences (mean ± SD across six exosome sources was 0.00825±0.02 miRNA molecules/exosome).^6^

Traditional methods such as quantitative reverse transcription polymerase chain reaction (qRT-PCR) for miRNA have been considered as the gold standard for miRNA quantification with femtomolar limits of detection.^7^ However, qRT-PCR requires complicated enzymatic amplification and complicated primer designs.^8^ Other quantification methods such as Northern blots^9^ and oligonucleotide microarrays^10–11^ are performed on cell lysate; Where fluorescent reporters^12–13^ are constructed to obtain enhanced fluorescence signals for profiling miRNA in cells.^14–17^ Traditional methods for miRNA detection do not meet current demands. In particular, loop-mediated isothermal amplification (LAMP),^18^ strand displacement amplification (SDA),^19^ exponential amplification reaction (EXPAR),^20–21^ rolling circle amplification (RCA),^22^ and some enzyme-free amplification methods, such as catalytic hairpin assembly (CHA),^23^ hybridization chain reaction (HCR),^24–25^ and entropy-driven catalysis^26^ have been employed widely for the high sensitivity detection of miRNA. Constrained by the detection limit and selectivity, none of these methods have been adopted for clinical use for exosomal miRNA detection without enzymatic target amplification. Therefore, there is an unmet need to develop an ultrasensitive and highly selective diagnostic approach without enzymatic amplification to effectively detect and quantify exosomal miRNAs.

Recently, our group developed a technique called Photonic Resonator Absorption Microscopy (PRAM) that can visualize individual gold nanoparticle (AuNPs) tags on a photonic crystal (PC) surface through resonance coupling.^27^ As described in prior publications,^27–33^ the detection principle of PRAM utilizes the resonant PC reflection at a wavelength of λ = 625 nm to provide a high reflected intensity from collimated low intensity LED illumination of the same wavelength into a webcam-variety image sensor. The AuNPs are strategically selected to provide strong absorption by localized surface plasmon resonance at the same wavelength.^34–36^ Thus, each surface-bound AuNP registers in the PC reflected image as a location with reduced intensity, compared to the surrounding regions without AuNPs. By immobilizing target-activated AuNP probes on a PC surface, PRAM has been used to quantify nucleic acids and proteins with single-particle resolution.^29–33^ As proof-of-concept studies for sensing miRNA, previous work focused on detecting chemically synthetic miRNAs, in which each detected miRNA molecule was associated with one AuNP tag^28, 31^ and lacked an amplification mechanism that could further reduce detection limits, as the target miRNA molecule is consumed by the detection process. To address this issue, we employ DNA-fueled molecular machines, including a series of toehold-mediated DNA strand displacement reactions (SDRs) involving target recycling^37^ as versatile tools for building switchable nanodevices,^38^ controlled nanoparticle assembly,^39–40^ mediated gene expression,^41^ and programmed DNA computation.^42^ In this work, we demonstrate substantial (> 400 folds) reduction of miRNA detection limits into sub-attomolar concentrations (0.24 aM) while decreasing the assay time to 20 minutes using PRAM detection in conjunction with target recycling by a DNA-fueled molecular machine. This assay is a single step, room temperature, one-pot reaction that requires only inexpensive synthetic nucleic acids and a single sample can be tested for less than five dollars. The TRAP method is also capable of multiplexing, with very low sample volume requirements (< 20 μL) and has potential for applications in frequent patient monitoring. Additionally, this assay has been applied to the detection of biological exosomal miRNAs and results have been validated with qRT-PCR.

## Results and Discussion

To demonstrate our TRAP method for miRNA detection, we choose a miRNA relevant to prostate cancer, miR-375 as the initial target.^43–44^ The miRNAs can be extracted from exosomes isolated from cell culture media (Scheme 1a) and the PC surface is prepared with an immobilized capture DNA (yellow) through a silanization process and then pretreated with a linker-protector complex consisting of a protector DNA (blue) and a linker DNA (green) that can link the capture DNA and protector DNAs through hybridizing both capture and protector DNAs. An important component of the linker-protector complex design is to leave free unhybridized regions at both 5’ and 3’ ends of the linker strand, resulting in toehold-1 and toehold-2, miRNA and probe DNA binding, respectively. All solutions are added to a reservoir made of Polydimethylsiloxane (PDMS) attached to the PC surface. Once the exosomal extraction samples are added into reaction wells, the miRNA target (red) binds to the free toehold-1 region on the linker strand (green) and replaces the protector strands (blue) through a DNA strand displacement reaction. Such a displacement of the protector strand by the miRNA target resulted in exposing the terminal segment (toehold-2) of the linker strand, which allows the probe DNA (pink) conjugated to AuNPs to invade the toehold-2 on linker strand and release the target miRNA with a second DNA strand displacement reaction. The tethered AuNPs can then be subsequently imaged by the reduction of reflected light intensity from the PC surface (Scheme 1a, 1b, 1c and Figure S2) in each location where an AuNP has bound. In our platform, the bound AuNPs on the surface can be enumerated using an automated image processing algorithm that identifies image pixels with reduced reflected intensity compared to the background, which is completed using a MATLAB script. Since the released target miRNA can then bind additional AuNPs, a Target Recycling Amplification Process (TRAP) was achieved by this design on the PRAM system, resulting in an increased nanoparticle binding and amplified signal.

**Scheme 1.**
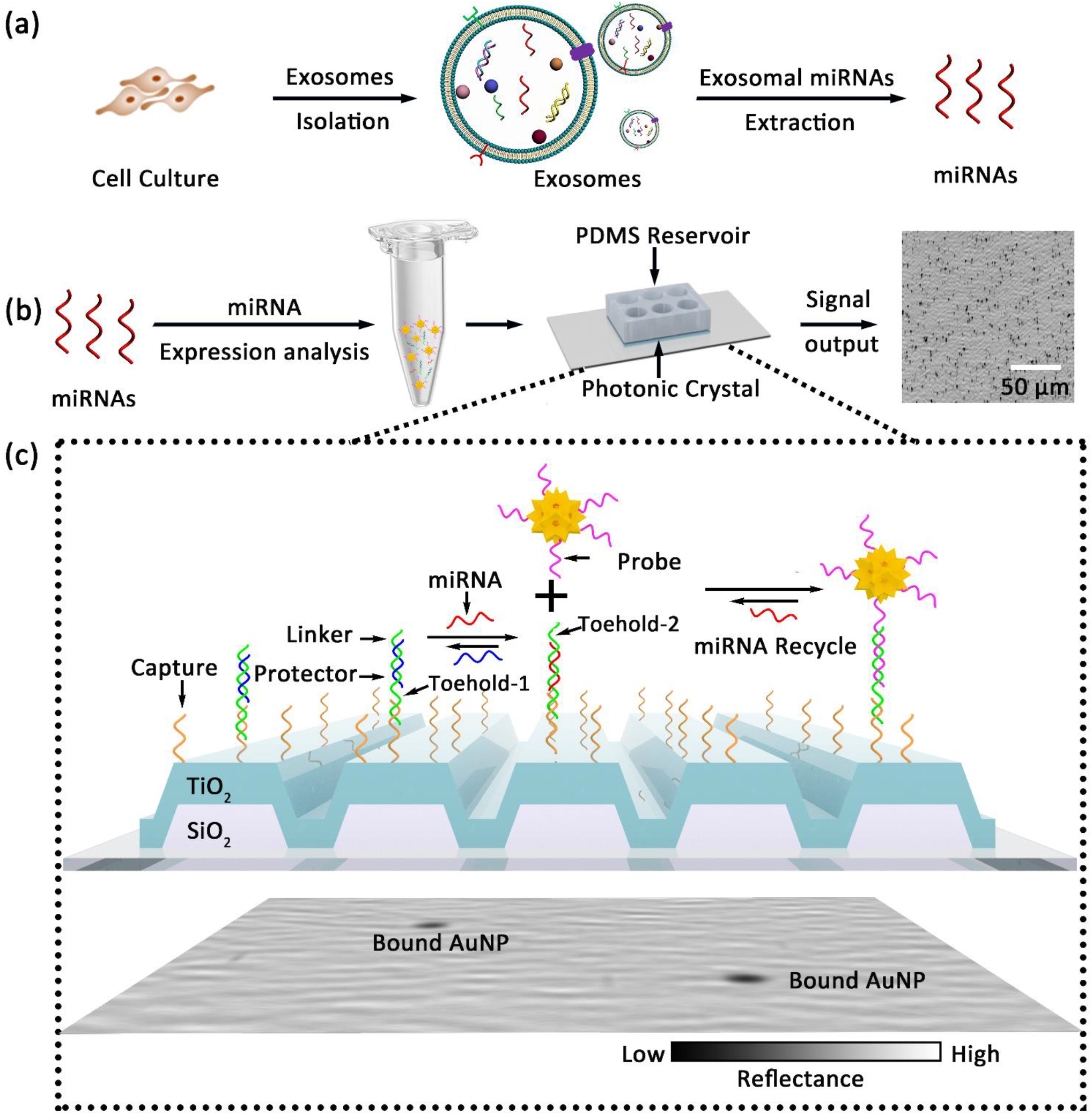
Schematic of miRNA detection workflow. (a) Extracted exosomal miRNAs are placed in a PDMS reservoir applied to the PC surface along with probes linked to AuNPs. (b) TRAP for digital resolution detection of miRNAs on a PC biosensor surface in which target miRNA displaces a protector strand on the PC-immobilized capture molecule to reveal a linker sequence by a strand displacement reaction. (c) DNA linker is pre-annealed with a partially complementary protector and hybridizes with DNA capture. In the presence the target miRNA, linker-protectors are “activated” through displacement of the protector, resulting in the exposure of an additional DNA linker sequence (toehold-2). Next, the probe sequence attached to the AuNP displaces the miRNA target from the capture molecule to simultaneously bind the AuNP to the PC surface, thus releasing the miRNA and making it available for another reaction. The PC-captured AuNPs are resolvable at single-particle resolution with high contrast through the synergistic coupling between the LSPR of AuNPs and the PC resonance.

To investigate the coupling behavior of the AuNPs with the PC, we performed a finite element method (FEM) simulation to investigate the near-field intensity distribution of the PC bound AuNP. As shown in Figure S2a, ~10^4^ field enhancement at the AuNP sharp tip features leads to enhanced absorption on the AuNP, which is consistent with previous results.^31^ The optical absorption of PC-coupled AuNPs was further analyzed by measuring the PC resonant reflected spectrum (Figure S2b, S2c). AuNP binding results in a localized quenching of the PC reflection intensity ΔI/I of approximately 12%, which is used as the contrast mechanism for image-based detection of single PC-attached nanoparticles.^45^

To achieve ultrasensitive limits of detection using the TRAP system, it is necessary to minimize non-specific AuNP binding and to demonstrate enhanced target-triggered signaling. In our TRAP design (Scheme 1), toehold-1 participates in the target miRNA-triggered strand displacement reaction, while toehold-2 allows for invasion of the probe-functionalized AuNP. Toehold-1 was designed with 5 bases according to the principles of nucleic acids involved in strand displacement reactions.^46^ Theoretically, a longer toehold-2 will result in faster kinetics and recycling of the target miRNAs.^47^ However, with a constant protector strand length, the longer toehold-2 will introduce more uncovered bases on the linker strand terminal and may cause non-specific binding in the absence of target miRNA. To optimize the design, four sets of linker sequences with different toehold-2 lengths were investigated. The binding characteristics of the different linker sequences were first analyzed with native PAGE (Figure S3). The four linkers with different sequence options were tested to ensure they would form stable capture-linker-protector structures and to test that the capture-linker-probe (C-L-P) complex only in the presence of the miRNA target. If this C-L-P is present in the absence of the miRNA target (Lanes 2, 5, 8, and 11), it would indicate that the length of toehold-2 allows for non-specific binding. The results for the longest linker sequence (L4 with a 4-nt initial toehold) suggest that the C-L-P formed even when no target was present, likely resulting from non-specific reactivity. In contrast, the shortest linker sequence (L1 with 1-nt initial toehold) did not form the C-L-P when the miRNA target was added, indicating that the target could not remove the protector strand. As shown in Figure S3, the linker strands (L2 and L3) containing 2 or 3 uncovered bases in the toehold-2 region produced a distinct C-L-P in the presence of the target but not in the absence of the target strand produce. To further optimize the effect of toehold length of the linker strand on results in PRAM imaging, we analyzed the L2, L3 and L4 with two, three, and four initial uncovered bases in the toehold-2 region, respectively. As shown in Figure S4, when the target miRNA is absent, the longest linker strand results in exceedingly high background signal that is consistent with the results from the PAGE experiments in Figure S3. However, after addition of a 10 pM concentration of target miRNA, the linker strand with only two initial uncovered bases on the toehold-2 region showed enhanced signal-to-noise of 142, which is 120-fold of that the linker with three free bases tested. Therefore, the design of two free bases for the initial uncovered linker terminal toehold (L2) was selected and used in all subsequent experiments. Kinetic studies showed that after addition of a 1 pM concentration of target miRNA the particle counts were saturated in 30 minutes (Figure S5).

In TRAP system, the DNA-probe functionalized AuNPs and PC tethered capture strands are bridged by the linker strand. Therefore the concentrations of the linker strands ranging from 0 to 50 pM were evaluated, and the bound AuNPs on the PC surface were counted (Figure S6). The captured AuNPs gradually increased from 133 to 491 when the linker concentrations were increased from 1 pM to 20 pM. However, the AuNPs on the PC surface decreased to 294 as the linker concentration was raised to 50 pM. If excess linker strands are present, both probe DNA and capture strands can hybridize with linker strands individually, resulting in fewer probe DNA and capture strands that are connected by the linker strands. This phenomenon was also demonstrated in the field of DNA guided gold nanoparticle assembly.^48^ Therefore, an optimal 20 pM concentration of the linker-protector complex is used in the following studies, with the protector in two times excess.

To find out the sensitivity of TRAP, we measured the dose-dependent immobilized AuNPs on the PC surface under various concentrations of target miRNA-375. In these experiments, a constant reaction volume of 20 μL solution, including target miRNA, the linker-protector complex, and probe DNA-modified AuNPs, was added into a PDMS reservoir. Separate wells were used for different concentrations of miRNAs ranging from 0.1 aM to 1 pM. Subsequently, the reactions were performed at room temperature and imaged after 10 min and 20 min. We observed an increased number of AuNPs on PC surface from 32, 82, 276, to 538 with higher concentrations of miRNA-375 from 0 aM, 1 aM, 1 fM to 1pM at 10 min (Figure 1a, additional PRAM images are shown in Figure S7). A linear calibration curve was calculated (Figure 1b) between 0.1 aM and 1 pM for miRNA-375 (R2 = 0.957) at 10 minutes. With a longer reaction time of 20 minutes (R2 = 0.999), the tethered AuNPs on PC increased by approximately 1.5-fold (Figure 1, and Figure S7) and the linear relationship between the particle counts and miRNA-375 concentrations also increases. A 20-minute reaction time provided a higher density of captured AuNPs, slightly better sensitivity and better signal-to-noise for miR-375. On this basis, the limit of detection (LOD), defined as the concentration at a signal threshold of three standard deviations (3σ) above background noise (3σ+blank), is calculated as 0.24 aM for miRNA-375 at 20 minutes. To demonstrate the robustness and universality of TRAP nucleic acid design, another target miRNA-21 was investigated and a LOD of 0.356 aM was observed (Figure S8).

**Figure 1.**
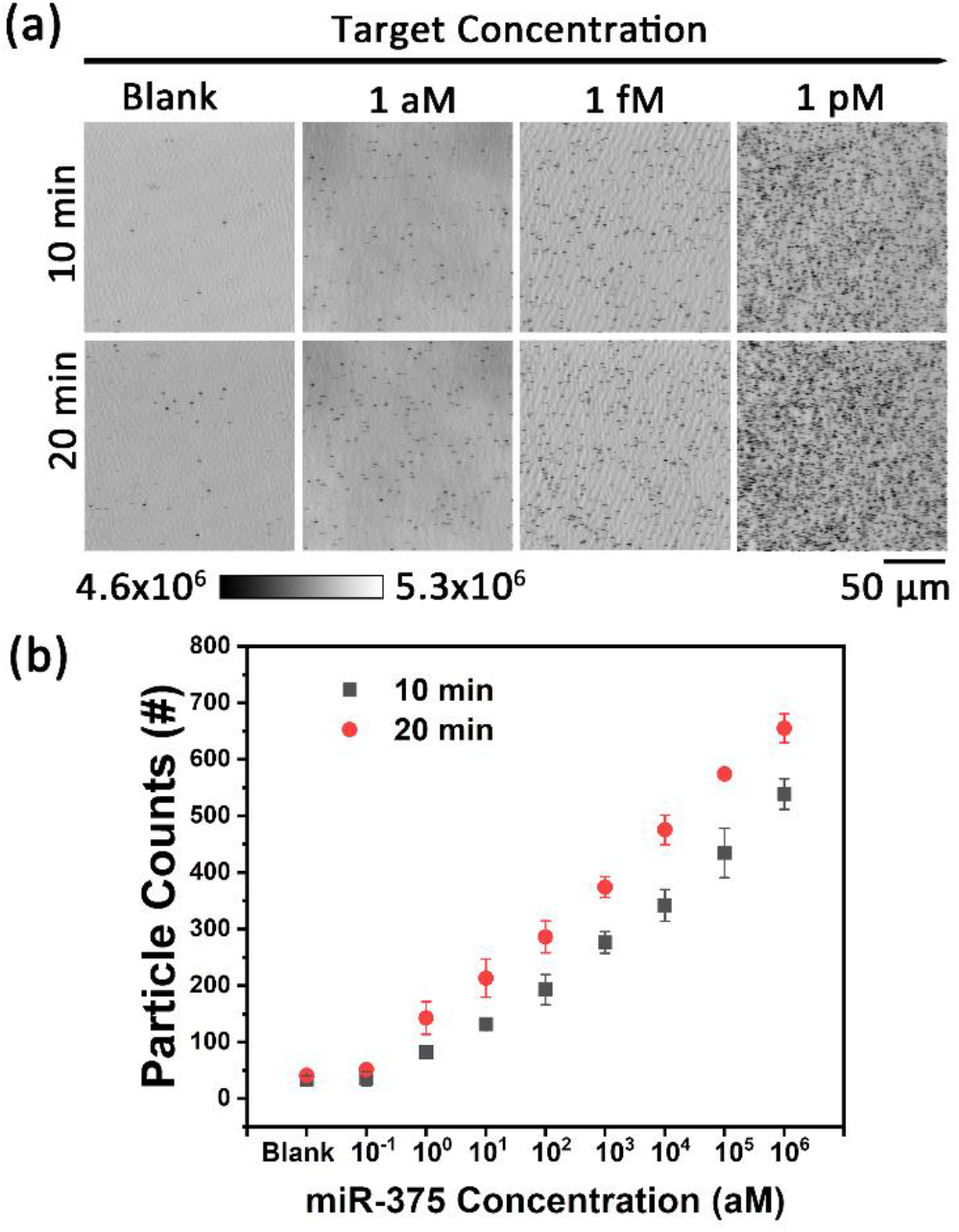
Kinetic discrimination of miRNA-375 using TRAP. (a) Dose-response TRAP images at single particle resolution at 10 mins and 20 mins. (b) Quantification of particle counts as a function of miRNA-375 concentration for three trials with standard error shown. Blank represents reaction without miRNA target present.

To examine the selectivity of TRAP, five different single nucleotide variants (SNVs) of miRNA-375, with the mismatch position at the 1st, 5th, 12th, 18th and 22nd from the 5’ end were tested in TRAP. The wild type of miRNA-375 target was tested at concentration of 1 fM, while the mismatched SNVs were tested at 1 pM. The TRAP images of wild type of miRNA-375 target showed ~302 nanoparticles while the mismatched SNVs have a background signal of 72 nanoparticles or fewer on PC (Figure 2). Even when the mismatch positions are located away from the initial toehold, and using a miRNA with a 1000-fold mismatch (1 pM) of the correct target, the resulting signal is still less than 25% of the signal of the correct target at 1 fM. Therefore, the TRAP system keeps high selectivity at single-base precision for miRNA detection. The branch migration reaction of the toehold strand displacement reaction additionally ensures selectivity, as a single base mismatch causes a Δ*G*° increase of +1.83 to +5.9 kcal/mol.^47^

**Figure 2.**
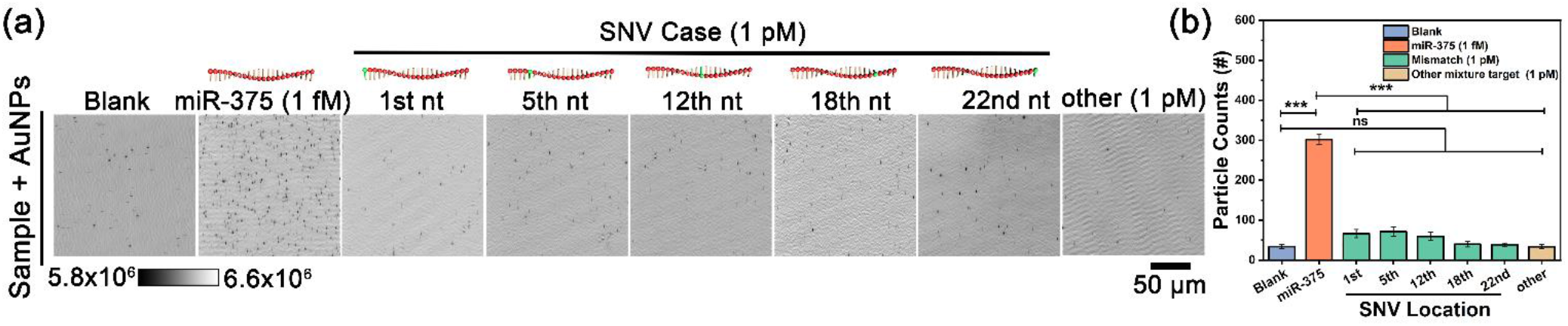
Selectivity of TRAP in detecting single nucleotide variants (SNVs) of miRNA-375. (a) TRAP images and (b) counted numbers of bound nanoparticle on PC surface in the presence of miRNA-375 or 5 different SNVs at the 1st, 5th, 12th, 18th and 22nd from 5’ terminal of miRNA. Approximately 6000:1 selectivity is demonstrated for detection of the target sequence against single base mismatched targets at each location investigated. (Two-tailed Mann-Whitney t test)

The TRAP system can be readily adapted to detect any miRNA sequence, and by measuring separate sub-volumes of a test sample in independent wells on the same PC biosensor, it is possible to perform multiple assays in parallel. To demonstrate this capability, three DNA strands were designed and optimized using the same principles and the feasibility of the DNA probes was confirmed with PAGE analysis (Figure S9). In the multiplex TRAP assays (Figure 3a), the same capture strands were used in every reservoir, and the only differences between the wells are the 1fM target miRNAs (Figure 3b), nanoparticle probes, and the specially designed linker-protector complexes. In the presence of 1 fM miRNAs, elevated counts of nanoparticles are observed upon the PC surface while the blank controls (with no target) displayed weak signals. All these results demonstrate that TRAP is applicable to a variety of miRNA targets that can be quantified simultaneously. To further verify the specificity of the TRAP assay, we used various controls including changing the linker-protector complex and the miRNAs to test the specificity. As shown in group 3 and group 4 in Figure S10, when using mismatched sensor sequences, using 1 pM of miRNA or other miRNA mixtures without the target, we do not observe a positive signal. On the other hand, in group 5, the use of the correct probe sequence and the corresponding very low concentration of target miRNA-21 (1fM) still produced a significant positive signal.

**Figure 3.**
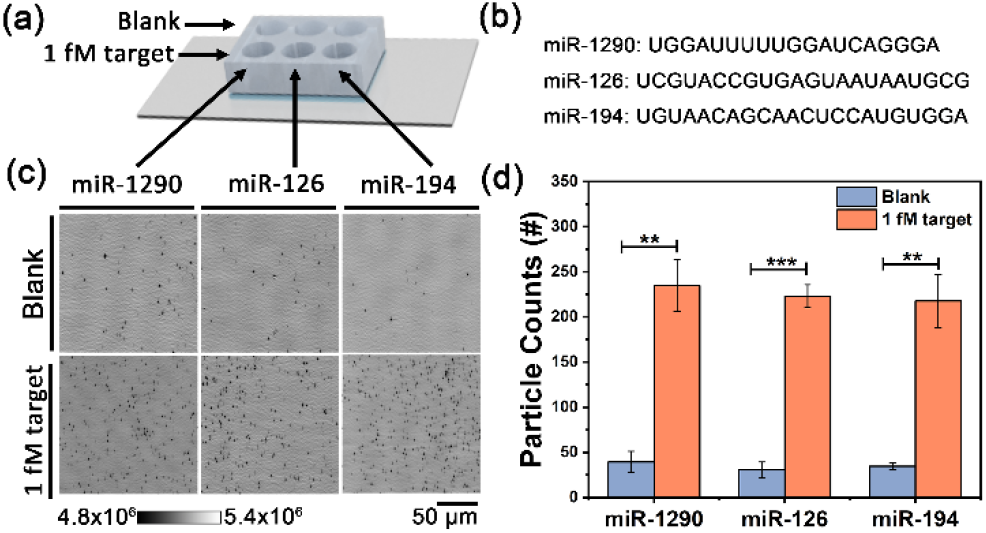
Multiplexed miRNA detection was performed in different reservoirs of one PC biosensor. (a) Utilization of a 6-well PDMS gasket applied to a single PC for simultaneous measurement of three negative controls and three separate miRNA targets; (b) The sequences of target miRNAs; (c) PRAM images and (d) particle counts on the PC surface with or without 1 fM miRNA targets. (Two-tailed Mann-Whitney t test)

MiRNA-375 and miRNA-21 have been identified as important biomarkers for prostate and breast cancer.^16, 43–44, 49–51^ The circulating miRNA-375 is significantly overexpressed in the blood of metastatic prostate cancer patients (compared to healthy individuals) and is involved in several processes affecting tumorigenesis and metastasis.^52–53^ As a proof-of-concept demonstration of TRAP on cancer diagnosis through monitoring the miRNAs expression in exosomes, miRNA-375 and miRNA-21 were chosen as the models. Total RNAs were first extracted from exosomes of a breast cancer cell line (MCF-7) and human prostate cancer cell line (DU145). Then, the extracted samples were used to test miRNA-375 and miRNA-21 expression in the TRAP system (Scheme 1). qRT-PCR quantification was simultaneously performed as a gold standard to validate the accuracy of TRAP (Figure S11). The detection limits of the TRAP approach for miRNA-375 and miRNA-21 in buffer are 0.15 copies/μL (0.24 aM of miRNA-375) and 0.21 copies/μL (0.356 aM of miRNA-21), which are 259-fold and 372-fold lower than that obtained by qRT-PCR for the same targets (Figure 4a). Through dilution of the exosomal miRNA extract, dose-responsive curves were obtained (Figure S12, S13 S14 and S15). The amount of miRNA-21 in MCF-7 cell-derived exosomes was higher (2535 times as much) than miRNA-375 and the amount of exosomal miRNA-21 in DU145 cell-derived exosomes was much higher (2412 times as much) than that of miRNA-375. At the same time, the amount of miRNA-375 and miRNA-21 in DU145 cell-derived exosomes was similar to that in MCF-7. We compare the miRNA concentration estimated by TRAP to that estimated using qRT-PCR, using calibration standards derived from multiple dilutions of the target. In Figure 4b, the results obtained by the two methods were consistent, which demonstrates the accuracy, reliability and practicability of the TRAP method. The detection limit of TRAP for miRNA-375 and miRNA-21 in cancer cell exosomes derived from MCF-7 and DU145 cultures are 1.2 copies/μL (2 aM of miRNA-375 and miRNA-21 in MCF-7 and DU145, respectively), which is 31-fold and 61-fold lower than that from qRT-PCR, respectively. The ultralow LOD will enable TRAP to be utilized for monitoring low-abundance miRNA in exosomes, especially for the down-regulated species encountered in cancer diagnosis. To demonstrate the feasibility of the TRAP digital sensing in clinical settings, miRNA-375 target was spiked into crude human plasma and serum at 1fM target concentration without any extraction and purification steps. The apparent positive signal in Figure S16 demonstrates that the TRAP method can quantify miRNA-375 directly from highly complex human plasma and serum samples with high sensitivity, offering the potential for rapid POC diagnostic testing of freely circulating miRNA biomarkers.

**Figure 4.**
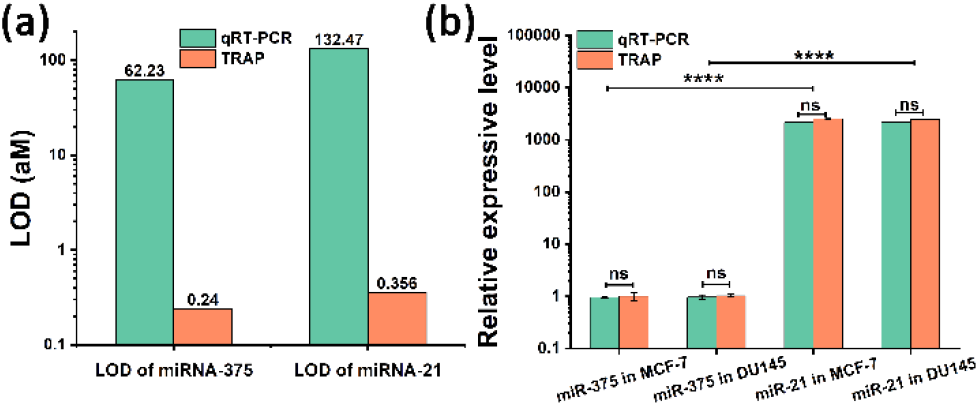
(a) Comparison of the limit of detection for exosomal miRNA-375 and miRNA-21 using TRAP and qRT-PCR methods. (b) Relative expressive levels of exosomal miRNA-375 and miRNA-21 derived from MCF-7 and DU145 cells were resolved with TRAP and qRT-PCR. The expressive level of miRNA-375 in MCF-7 cells was defined as 1. The relative expressive level was compared to miRNA-375 in MCF-7 cells. (calculated by an unpaired, two-tailed t test)

## Conclusion

In conclusion, a target recycling amplification process (TRAP) detection method was developed using photonic resonator absorption microscopy, which shows exciting potential for monitoring exosomal miRNAs with ultra-sensitivity and single-base mismatch selectivity. The approach is a single-step, wash-free, enzyme-free, isothermal, 20-minute, room temperature process that can be readily adapted toward any miRNA target through different DNA probe designs. The AuNP-probe complex is designed to be universally used for all targets. The TRAP method achieves cancer biomarker miRNA detection with a detection limit of 0.24 aM for miRNA-375 and 0.356 aM of miRNA-21. The target miRNA concentration can be measured over a broad range from 1 aM to 1 pM with single-base precision. Additionally, TRAP can perform multiplexed miRNA detection in a one-batch manner with multiplexing, consistent performance. Compared with traditional qRT-PCR methods, TRAP showed similar accuracy in profiling exosomal miRNAs derived from cancer cells, but also exhibited at least 31-fold and 61-fold enhancement in the limits of miRNA-375 and miRNA-21 detection, respectively. These features show promise for early detection of cancer through monitoring changes in exosomal miRNAs or other circulating nucleic acid biomarkers and demonstrates a simple approach compatible with point of care scenarios for longitudinally measuring therapy effectiveness and monitoring remission.

## Supporting information

Essential Experimental Procedures and Data

## Acknowledgements

This work was supported by National Institutes of Health (R01EB029805). M.K. was also supported by National Institutes of Health (R01-CA212097). N.L. acknowledges support from Zhejiang University ZJU-UIUC Joint Research Center (DREMES202001) and from the Mikashi award of the Carl R. Woese Institute for Genomic Biology. B. Z. acknowledges support from Genomic Diagnostics at the Woesse Institute for Genomic Biology and the Grainger College of Engineering. Y.X. acknowledges support from the Cancer Center at Illinois C*STAR program. We thank Dr. Hui Xu and Huimin Zhang (Tumor Engineering and Phenotyping Facility, Cancer Center at Illinois, UIUC) for help with the experiments to extract exosomal miRNA from cell culture and qPCR quantification.

## Data Availability Statement

Essential Experimental Procedures/Data. The data that support the findings of this study are available in the Supporting Information of this article.

## References

(1) Cowland, J. B.; Hother, C.; Grønbaek, K., MicroRNAs and cancer. APMIS 2007, 115, 1090–106.

(2) Alexander, M.; Hu, R.; Runtsch, M. C.; Kagele, D. A.; Mosbruger, T. L.; Tolmachova, T.; Seabra, M. C.; Round, J. L.; Ward, D. M.; O’Connell, R. M., Exosome-delivered microRNAs modulate the inflammatory response to endotoxin. Nat. Commun. 2015, 6, 7321.

(3) Zheng, D.; Huo, M.; Li, B.; Wang, W.; Piao, H.; Wang, Y.; Zhu, Z.; Li, D.; Wang, T.; Liu, K., The Role of Exosomes and Exosomal MicroRNA in Cardiovascular Disease. Front. Cell Dev. Biol. 2021, 8.

(4) Nedaeinia, R.; Manian, M.; Jazayeri, M. H.; Ranjbar, M.; Salehi, R.; Sharifi, M.; Mohaghegh, F.; Goli, M.; Jahednia, S. H.; Avan, A.; Ghayour-Mobarhan, M., Circulating exosomes and exosomal microRNAs as biomarkers in gastrointestinal cancer. Cancer Gene Ther. 2017, 24, 48–56.

(5) Köberle, V.; Pleli, T.; Schmithals, C.; Augusto Alonso, E.; Haupenthal, J.; Bönig, H.; Peveling-Oberhag, J.; Biondi, R. M.; Zeuzem, S.; Kronenberger, B.; Waidmann, O.; Piiper, A., Differential stability of cell-free circulating microRNAs: implications for their utilization as biomarkers. PLoS One 2013, 8, e75184.

(6) Chevillet, J. R.; Kang, Q.; Ruf, I. K.; Briggs, H. A.; Vojtech, L. N.; Hughes, S. M.; Cheng, H. H.; Arroyo, J. D.; Meredith, E. K.; Gallichotte, E. N.; Pogosova-Agadjanyan, E. L.; Morrissey, C.; Stirewalt, D. L.; Hladik, F.; Yu, E. Y.; Higano, C. S.; Tewari, M., Quantitative and stoichiometric analysis of the microRNA content of exosomes. Proc. Natl. Acad. Sci. U. S. A. 2014, 111, 14888–14893.

(7) Causa, F.; Aliberti, A.; Cusano, A. M.; Battista, E.; Netti, P. A., Supramolecular Spectrally Encoded Microgels with Double Strand Probes for Absolute and Direct miRNA Fluorescence Detection at High Sensitivity. J. Am. Chem. Soc. 2015, 137, 1758–1761.

(8) Moltzahn, F.; Olshen, A. B.; Baehner, L.; Peek, A.; Fong, L.; Stöppler, H.; Simko, J.; Hilton, J. F.; Carroll, P.; Blelloch, R., Microfluidic-based multiplex qRT-PCR identifies diagnostic and prognostic microRNA signatures in the sera of prostate cancer patients. Cancer Res. 2011, 71, 550–60.

(9) Lee, E. J.; Baek, M.; Gusev, Y.; Brackett, D. J.; Nuovo, G. J.; Schmittgen, T. D., Systematic evaluation of microRNA processing patterns in tissues, cell lines, and tumors. RNA 2008, 14, 35–42.

(10) Pritchard, C. C.; Cheng, H. H.; Tewari, M., MicroRNA profiling: approaches and considerations. Nat. Rev. Genet. 2012, 13, 358–69.

(11) Fabri-Faja, N.; Calvo-Lozano, O.; Dey, P.; Terborg, R. A.; Estevez, M. C.; Belushkin, A.; Yesilköy, F.; Duempelmann, L.; Altug, H.; Pruneri, V.; Lechuga, L. M., Early sepsis diagnosis via protein and miRNA biomarkers using a novel point-of-care photonic biosensor. Anal. Chim. Acta 2019, 1077, 232–242.

(12) Li, W.; Jiang, W.; Ding, Y.; Wang, L., Highly selective and sensitive detection of miRNA based on toehold-mediated strand displacement reaction and DNA tetrahedron substrate. Biosens. Bioelectron. 2015, 71, 401–406.

(13) Kim, D.; Wei, Q.; Kim, D. H.; Tseng, D.; Zhang, J.; Pan, E.; Garner, O.; Ozcan, A.; Di Carlo, D., Enzyme-Free Nucleic Acid Amplification Assay Using a Cellphone-Based Well Plate Fluorescence Reader. Anal. Chem. 2018, 90, 690–695.

(14) Deng, R.; Tang, L.; Tian, Q.; Wang, Y.; Lin, L.; Li, J., Toehold-initiated rolling circle amplification for visualizing individual microRNAs in situ in single cells. Angew. Chem., Int. Ed. 2014, 53, 2389–93.

(15) Hemphill, J.; Deiters, A., DNA computation in mammalian cells: microRNA logic operations. J. Am. Chem. Soc. 2013, 135, 10512–8.

(16) Wang, X.; Yan, N.; Song, T.; Wang, B.; Wei, B.; Lin, L.; Chen, X.; Tian, H.; Liang, H., Robust Fuel Catalyzed DNA Molecular Machine for in Vivo MicroRNA Detection. Adv. Biosyst. 2017, 1, 1700060–n/a.

(17) Yan, N.; Wang, X. J.; Lin, L.; Song, T. J.; Sun, P. J.; Tian, H. Y.; Liang, H. J.; Chen, X. S., Gold Nanorods Electrostatically Binding Nucleic Acid Probe for In Vivo MicroRNA Amplified Detection and Photoacoustic Imaging-Guided Photothermal Therapy. Adv. Funct. Mater. 2018, 28.

(18) Li, C.; Li, Z.; Jia, H.; Yan, J., One-step ultrasensitive detection of microRNAs with loop-mediated isothermal amplification (LAMP). Chem. Commun. 2011, 47, 2595–2597.

(19) Walker, G. T.; Fraiser, M. S.; Schram, J. L.; Little, M. C.; Nadeau, J. G.; Malinowski, D. P., Strand displacement amplification--an isothermal, in vitro DNA amplification technique. Nucleic Acids Res 1992, 20, 1691–6.

(20) Yang, L.; Fang, J.; Li, J.; Ou, X.; Zhang, L.; Wang, Y.; Weng, Z.; Xie, G., An integrated fluorescence biosensor for microRNA detection based on exponential amplification reaction-triggered three-dimensional bipedal DNA walkers. Anal. Chim. Acta 2021, 1143, 157–165.

(21) Li, R.-D.; Yin, B.-C.; Ye, B.-C., Ultrasensitive, colorimetric detection of microRNAs based on isothermal exponential amplification reaction-assisted gold nanoparticle amplification. Biosens. Bioelectron. 2016, 86, 1011–1016.

(22) Ali, M. M.; Li, F.; Zhang, Z.; Zhang, K.; Kang, D.-K.; Ankrum, J. A.; Le, X. C.; Zhao, W., Rolling circle amplification: a versatile tool for chemical biology, materials science and medicine. Chem. Soc. Rev. 2014, 43, 3324–3341.

(23) Bao, J.; Hou, C.; Zhao, Y.; Geng, X.; Samalo, M.; Yang, H.; Bian, M.; Huo, D., An enzyme-free sensitive electrochemical microRNA-16 biosensor by applying a multiple signal amplification strategy based on Au/PPy–rGO nanocomposite as a substrate. Talanta 2019, 196, 329–336.

(24) Wu, Z.; Liu, G.-Q.; Yang, X.-L.; Jiang, J.-H., Electrostatic Nucleic Acid Nanoassembly Enables Hybridization Chain Reaction in Living Cells for Ultrasensitive mRNA Imaging. J. Am. Chem. Soc. 2015.

(25) Nie, Y.; Yuan, X.; Zhang, P.; Chai, Y. Q.; Yuan, R., Versatile and Ultrasensitive Electrochemiluminescence Biosensor for Biomarker Detection Based on Nonenzymatic Amplification and Aptamer-Triggered Emitter Release. Anal. Chem. 2019, 91, 3452–3458.

(26) Liang, C.-P.; Ma, P.-Q.; Liu, H.; Guo, X.; Yin, B.-C.; Ye, B.-C., Rational Engineering of a Dynamic, Entropy-Driven DNA Nanomachine for Intracellular MicroRNA Imaging. Angew. Chem., Int. Ed. 2017, 56, 9077–9081.

(27) Zhuo, Y.; Hu, H.; Chen, W.; Lu, M.; Tian, L.; Yu, H.; Long, K. D.; Chow, E.; King, W. P.; Singamaneni, S.; Cunningham, B. T., Single nanoparticle detection using photonic crystal enhanced microscopy. Analyst 2014, 139, 1007–1015.

(28) Che, C.; Xue, R.; Li, N.; Gupta, P.; Wang, X.; Zhao, B.; Singamaneni, S.; Nie, S.; Cunningham, B. T., Accelerated Digital Biodetection Using Magneto-plasmonic Nanoparticle-Coupled Photonic Resonator Absorption Microscopy. ACS Nano 2022, 16, 2345–2354.

(29) Ghosh, S.; Li, N.; Xiong, Y.; Ju, Y.-G.; Rathslag, M. P.; Onal, E. G.; Falkiewicz, E.; Kohli, M.; Cunningham, B. T., A compact photonic resonator absorption microscope for point of care digital resolution nucleic acid molecular diagnostics. Biomed. Opt. Express 2021, 12, 4637–4650.

(30) Che, C.; Li, N.; Long, K. D.; Aguirre, M. Á.; Canady, T. D.; Huang, Q.; Demirci, U.; Cunningham, B. T., Activate capture and digital counting (AC + DC) assay for protein biomarker detection integrated with a self-powered microfluidic cartridge. Lab on a Chip 2019, 19, 3943–3953.

(31) Canady, T. D.; Li, N.; Smith, L. D.; Lu, Y.; Kohli, M.; Smith, A. M.; Cunningham, B. T., Digital-resolution detection of microRNA with single-base selectivity by photonic resonator absorption microscopy. Proc. Natl. Acad. Sci. U. S. A. 2019, 116, 19362–19367.

(32) Zhao, B.; Che, C.; Wang, W.; Li, N.; Cunningham, B. T., Single-step, wash-free digital immunoassay for rapid quantitative analysis of serological antibody against SARS-CoV-2 by photonic resonator absorption microscopy. Talanta 2021, 225, 122004.

(33) Zhao, B.; Wang, W.; Li, N.; Garcia-Lezana, T.; Che, C.; Wang, X.; Losic, B.; Villanueva, A.; Cunningham, B. T., Digital-resolution and highly sensitive detection of multiple exosomal small RNAs by DNA toehold probe-based photonic resonator absorption microscopy. Talanta 2022, 241, 123256.

(34) Draz, M. S.; Shafiee, H., Applications of gold nanoparticles in virus detection. Theranostics 2018, 8, 1985–2017.

(35) Peng, H.-I.; Miller, B. L., Recent advancements in optical DNA biosensors: Exploiting the plasmonic effects of metal nanoparticles. Analyst 2011, 136, 436–447.

(36) Altug, H.; Oh, S.-H.; Maier, S. A.; Homola, J., Advances and applications of nanophotonic biosensors. Nat. Nanotechnol. 2022, 17, 5–16.

(37) Yurke, B.; Turberfield, A. J.; Mills, A. P.; Simmel, F. C.; Neumann, J. L., A DNA-fuelled molecular machine made of DNA. Nature 2000, 406, 605–608.

(38) Jung, C.; Allen, P. B.; Ellington, A. D., A stochastic DNA walker that traverses a microparticle surface. Nat. Nanotechnol. 2016, 11, 157–163.

(39) Song, T.; Xiao, S.; Yao, D.; Huang, F.; Hu, M.; Liang, H., An Efficient DNA-Fueled Molecular Machine for the Discrimination of Single-Base Changes. Adv. Mater. 2014, 26, 6181–6185.

(40) Song, T.; Liang, H., Synchronized Assembly of Gold Nanoparticles Driven by a Dynamic DNA-Fueled Molecular Machine. J. Am. Chem. Soc. 2012, 134, 10803–10806.

(41) Green, Alexander A.; Silver, Pamela A.; Collins, James J.; Yin, P., Toehold Switches: De-Novo-Designed Regulators of Gene Expression. Cell 2014, 159, 925–939.

(42) Qian, L.; Winfree, E., Scaling Up Digital Circuit Computation with DNA Strand Displacement Cascades. Science 2011, 332, 1196–1201.

(43) Wang, Y.; Lieberman, R.; Pan, J.; Zhang, Q.; Du, M.; Zhang, P.; Nevalainen, M.; Kohli, M.; Shenoy, N. K.; Meng, H.; You, M.; Wang, L., miR-375 induces docetaxel resistance in prostate cancer by targeting SEC23A and YAP1. Mol. Cancer 2016, 15, 70.

(44) Huang, X.; Yuan, T.; Liang, M.; Du, M.; Xia, S.; Dittmar, R.; Wang, D.; See, W.; Costello, B. A.; Quevedo, F.; Tan, W.; Nandy, D.; Bevan, G. H.; Longenbach, S.; Sun, Z.; Lu, Y.; Wang, T.; Thibodeau, S. N.; Boardman, L.; Kohli, M.; Wang, L., Exosomal miR-1290 and miR-375 as prognostic markers in castration-resistant prostate cancer. Eur. Urol. 2015, 67, 33–41.

(45) Huang, Q.; Canady, T. D.; Gupta, R.; Li, N.; Singamaneni, S.; Cunningham, B. T., Enhanced Plasmonic Photocatalysis through Synergistic Plasmonic–Photonic Hybridization. ACS Photonics 2020, 7, 1994–2001.

(46) Zhang, D. Y.; Winfree, E., Control of DNA Strand Displacement Kinetics Using Toehold Exchange. J. Am. Chem. Soc. 2009, 131, 17303–17314.

(47) Zhang, D. Y.; Chen, S. X.; Yin, P., Optimizing the specificity of nucleic acid hybridization. Nat. Chem. 2012, 4, 208–214.

(48) Mirkin, C. A.; Letsinger, R. L.; Mucic, R. C.; Storhoff, J. J., A DNA-based method for rationally assembling nanoparticles into macroscopic materials. Nature 1996, 382, 607–609.

(49) Ward, A.; Balwierz, A.; Zhang, J. D.; Küblbeck, M.; Pawitan, Y.; Hielscher, T.; Wiemann, S.; Sahin, Ö., Re-expression of microRNA-375 reverses both tamoxifen resistance and accompanying EMT-like properties in breast cancer. Oncogene 2013, 32, 1173–82.

(50) Zou, L.; Wu, Z.; Liu, X.; Zheng, Y.; Mei, W.; Wang, Q.; Yang, X.; Wang, K., DNA Hydrogelation-Enhanced Imaging Ellipsometry for Sensing Exosomal microRNAs with a Tunable Detection Range. Anal. Chem. 2020, 92, 11953–11959.

(51) Zhao, J.; Liu, C.; Li, Y.; Ma, Y.; Deng, J.; Li, L.; Sun, J., Thermophoretic Detection of Exosomal microRNAs by Nanoflares. J. Am. Chem. Soc. 2020, 142, 4996–5001.

(52) Brase, J. C.; Johannes, M.; Schlomm, T.; Fälth, M.; Haese, A.; Steuber, T.; Beissbarth, T.; Kuner, R.; Sültmann, H., Circulating miRNAs are correlated with tumor progression in prostate cancer. Int. J. Cancer 2011, 128, 608–16.

(53) Cai, S.; Pataillot-Meakin, T.; Shibakawa, A.; Ren, R.; Bevan, C. L.; Ladame, S.; Ivanov, A. P.; Edel, J. B., Singlemolecule amplification-free multiplexed detection of circulating microRNA cancer biomarkers from serum. Nat. Commun. 2021, 12, 3515.

