## Supplementary material for "Target Recycling Amplification Process for Digital Detection of Exosomal MicroRNAs Through Photonic Resonator Absorption Microscopy": Essential Experimental Procedures and Data

**Photonic crystals.** All the photonic crystals (PCs) used in experiments were obtained from Moxtek (Orem, UT). The PC structure is comprised of a low refractive index periodic grating structure that is coated with a higher refractive index material ($TiO_2$). PCs are fabricated on glass 8-inch diameter glass wafers deposited with a 10 nm etch stop layer of $Al_2O_3$. The periodic grating patterns are constructed by depositing a layer of $SiO_2$ followed by large area ultraviolet interference lithography. Finally, a thin layer of $TiO_2$ (thickness ~100 nm) is deposited on the etched wafers. The resulting PCs are diced into smaller chips of 1 × 1.2 $cm^2$. The PC is designed to function as a narrowband optical resonator that optimally reflects $\lambda$ = 625 nm with nearly 100% efficiency under water immersion.[1]

**Methods.**

**DNA sequence design:** Random sequences composed of only A, T, G, and C were designed with the help of NUPACK software to reduce secondary structures and interactions. The domain sequences were verified again with NUPACK to ensure the selected domain sequences possessed minimal secondary structure and crosstalk. For the alkanethiol modification sequences, a 10 poly-A spacer was chosen due to its known low interaction with the gold interface to improve sample stability and coverage. Toehold probes were designed to be robust for changes in temperature, concentration, and salinity by considering $\Delta\Delta\mathrm{G}° = \Delta G°(SC) - \Delta G°(XC) = 0$, where SC is spurious target-linker complex, and XC is the correct target according to Zhang et al. The branch migration reaction of the toehold strand displacement reaction additionally ensures selectivity, as a single base mismatch causes a $\Delta G°$ increase of +1.83 to +5.9 kcal/mol.[2] $\Delta G°$ values were also examined in NUPACK.

**Nucleic acids.** All the oligonucleotides used in the TRAP capture system were purchased from Integrated DNA Technologies (Coralville, Iowa), with standard purification. The concentrations of the oligonucleotides were calculated based on the molar extinction coefficient of single-stranded DNA. The same PC capture DNA sequence is used for all five DNA sequences. The probe sequence is terminated with a 5' dithiol group. The capture sequence is terminated with a 3' amine group. The sequences of the oligonucleotides used are shown below.

PC capture: AGTGAGAGTGGGATG AAAAAAAAAA/3AmMO

For miRNA-375:

MiRNA-375: rUrUrUrGrUrUrCrGrUrUrCrGr GrCrUrCrG rCrGrUrGrA

Target-miRNA375: TTT GTT CGT TCG GCT C GCGTGA

Linker-375: CATCCCACTCTCACTTCACGCGAGCCGAACGAACAAACAACTC

Protector-375: GTTGTTTGTTCGTTCGGCTCG

Probe-375: /5ThioMC6-D/ AAAAAAAAAA GGGAGTTGTTTGTTCGTTCGGCTCG

L1: CATCCCACTCTCACTTCACGCGAGCCGAACGAACAAACAACT

L2: CATCCCACTCTCACTTCACGCGAGCCGAACGAACAAACAACTC

L3: CATCCCACTCTCACTTCACGCGAGCCGAACGAACAAACAACTCC

L4: CATCCCACTCTCACTTCACGCGAGCCGAACGAACAAACAACTCCC

Mismatch miR-375 ($MM_x$, where x denotes mismatch position from 5' end)

$MM_1$: **C**TTGTTCGTTCG GCTCG CGTGA

$MM_5$: TTTG**A**TCGTTCG GCTCG CGTGA

$MM_{12}$: TTTGTTCGTTC**C** GCTCG CGTGA

$MM_{18}$: TTTGTTCGTTCG GCTCG **A**GTGA

$MM_{22}$: TTTGTTCGTTCG GCTCG CGTG**C**

For miRNA-1290:

MiRNA-1290: rUrGrGr ArUrU rUrUrU rGrGrA rUrCrA rGrGrG rA

Target-miRNA1290: TGG ATT TTT GGA TCA GGG A

Linker-1290: CATCCCACTCTCACT  TCCCT GATCCAAAAATCCA  CAAC TC

Protector-1290: GTTG TG GAT TTT TGG ATC

Probe-1290: /5ThioMC6-D/AAAAAAAAAA  GGGAGTTG  TGGATTTTTGGATC

For miRNA-21:

miRNA-21: rUrArG rCrUrU rArUrC rArGrA rCrUrG rArUrG rUrUrG rA

Target-miRNA21: TAG CTT ATC AGA CTG ATG TTG A

Linker-21: CATCCCACTCTCACT  TCAAC ATCAGTCTGATAAGCTA CAAC TC

Protector-21: GTTG TAGCTTATCAGACTGAT

Probe-21: /5ThioMC6-D/AAAAAAAAAA  GGGAGTTG TAGCTTATCAGACTGAT

For miRNA-194:

miRNA-194: rUrGrU rArArC rArGrC rArArC rUrCrC rArUrG rUrGrG rA

Target-miRNA194: TGT AAC AGC AAC TCC ATG TGG A

Linker-194: CATCCCACTCTCACT  TCCAC ATGGAGTTGCTGTTACA CGAA TC

Protector-194: TTCG TGTAACAGCAACTCCAT

Probe-194: /5ThioMC6-D/AAAAAAAAAA  GATTCG TGTAACAGCAACTCCAT

For miRNA-126:

miRNA-126: rUrCrG rUrArC rCrGrU rGrArG rUrArA rUrArA rUrGrC rG

Target-miRNA126: TCG TAC CGT GAG TAA TAA TGC G

Linker-126: CATCCCACTCTCACT  CGCAT TATTACTCACGGTACGA CAAC TC

Protector-126: GTTG TCGTACCGTGAGTAATA

Probe-126: /5ThioMC6-D/AAAAAAAAAA GGGAGTTG TCGTACCGTGAGTAATA

**Instrumentations.** The UV-vis spectra were obtained from a Thermo Scientific Nanodrop Onec Microvolume UV-vis Spectrophotometer W/ Cuvette.

**Preparation of double stranded linker-protector (LP).** The protector oligonucleotide strand was annealed to the linker sequence with a stoichiometric ratio of 1:2 with the protector in excess. For the annealing process, the oligos were heated to 95 °C for ten minutes then were allowed to cool to room temperature. The annealed linker-protector was diluted in a 1xTE, 12.5 mM $MgCl_2$ and 0.025% Tween20 buffer, then the product was stored at 4 °C until use.

**Polyacrylamide gel electrophoresis.** 12% polyacrylamide gel was employed for the verification of the formation of the TRAP nucleic acid complexes and products. All reaction products were loaded on the 1.5 mm thin gel. Electrophoresis was carried out at 165 V for 40 min at room temperature in 1×TBE buffer. After separation, the gel was stained by gel red and imaged with the Bio-red fluorescence gel imaging system.

**Nanoparticle conjugation**.

1mL 80-nm diameter NanoUrchin AuNPs (Cytodiagnostics, Inc.) were functionalized with thiol-modified DNA via gold-sulfur chemistry. DNA-AuNP complexes were prepared as previously described with modifications. Thiol-modified DNA was activated with two equivalents of tris(2-carboxyethyl) phosphine hydrochloride (TCEP). 80-nm NanoUrchin AuNPs were functionalized by mixing deprotected alkanethiol oligonucleotides with aqueous nanoparticle solution (particle concentration 1 OD) to a final probe strand concentration of 100 nM, then add m-PEG1K to a final concentration of 80 µg/mL. After ~48 h, to remove excess thiol-DNA, the solution was centrifuged (800 rcf, 10 min) and the supernatant was carefully removed using a micropipette. The deposited DNA-AuNPs were rinsed with an equal volume of 10 mM TE (0.025% Tween20, pH 7.4). The centrifuging/rinsing procedure was repeated two times. The final deposition was resuspended in stock solution (10 mM TE 0.025% Tween20, pH 7.4). and stored at 4°C for further use.

**Kinetic study.**

The assay was performed by mixing a constant amount of DNA-AuNP and a defined concentration of miRNA-375 in PC-PDMS wells. After miRNA-375 was added, the analysis was performed at 10-minute intervals for a total time 90 minutes immediately.

**Optimization of linker strand concentration.**

In our design, two different kinds of thiol-modified DNA strand (probe DNA) and amino-modified DNA (capture DNA) are conjugated on AuNPs and the PC surface. The linker strand acts like a bridge to connect the PC capture DNA and the probe DNA on the AuNPs. To optimize the appropriate concentration of linker chains in the system, the optimized ratio of linker strand with nanoparticle and capture on PC are performed. By adding different amounts of linker strand, the linker directed connect AuNPs on the PC surface is recorded by testing the PRAM imaging. As shown in Fig. S6, increased nanoparticle density in PRAM images are observed when the linker concentration increases from 0 to 20 pM. However, we observe a decreased nanoparticle density with the addition of 50 pM. These results indicate that under the conditions of probe DNA, capture DNA and 20 pM linker DNA ratios, high AuNPs can be immobilized on the PC surface within 90 minutes. In the next experiment, although a complex strand composed of linker and protector was used, the ratio of linker strand to AuNPs and capture-DNA was the same.

### Cell culture and exosomal total RNA isolation

MCF-7 and DU145 cells were cultured in Eagle's Minimum Essential Medium (EMEM) with 10% exosome-depleted fetal bovine serum (FBS, Gibco, catalog number A2720803). Cells were incubated at 37 °C in 5% $CO_2$ for 72 hours to 90% confluency. Exosomes from cell culture supernatant were isolated with Total Exosome Isolation Reagent (Invitrogen, catalog number 4478359). The reagent was added to the cell culture media, and the solution is incubated overnight at 2°C to 8°C. After centrifugation at 10,000 x g for 60 min, the pellet of exosomes is resuspended in PBS. Then total RNA was extracted from exosomes by using Total Exosome RNA & Protein Isolation Kit (Invitrogen, catalog number: 4478545).

### miRNA quantification by qPCR.

miR-21 and miR-375 were quantified by qPCR with Pre-designed TaqMan primers (Applied Biosystems, catalog number 4427975) for miR-21 and miR-375, respectively. Complementary DNAs (cDNAs) were synthesized by using TaqMan™ MicroRNA Reverse Transcription Kit (Applied Biosystems, catalog number 4366596). 10 ng of total RNA was added as described by protocol, reaction at 16℃ for 30mins, 42℃ for 30 mins and reaction stop at 85℃ for 5 mins. The cDNA product was diluted by 20-fold and 2 µl of diluted cDNA was mixed with TaqMan Universal Master Mix II (Applied Biosystems, catalog number 4440043), 1 µl of primers, and RNase-free water to a final volume of 20 µl. The reaction was performed and monitored by a real-time PCR detection system (QuantStudio™ 3 Real-Time PCR System) using the following thermocycler conditions: pre-denature at 95℃ for 10 min and 40 cycles of denature (94℃ for 40s) and anneal (60℃ for 30s). For absolute quantification of miR-375 and miR-21 in MCF-7 and DU 145, a standard curve was established using synthetic miR-375 AND miR-21 with different concentrations ranging from 1fM to 100 nM.

**Direct quantification of miR-375 in human serum and plasma.**

To demonstrate future clinical potential, RNA sequences for miR-375 were spiked into buffer, crude human serum, and human plasma at a concentration of 1 fM at room temperature, with no further purification steps. The TRAP pre-hybridized LP complex and DNA-AuNPs were added to the detection well, along with the spiked miRNA-375 samples and a negative control, then were imaged using the PRAM system after 15 minutes.

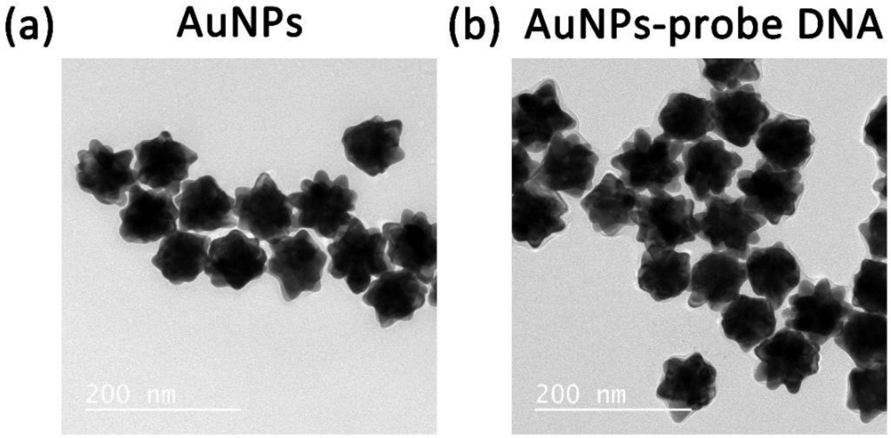


**Figure S1.** TEM imaging of NanoUrchin AuNPs (a), and conjugate product of NanoUrchin AuNPs with probe DNA and m-PEG1K (b).

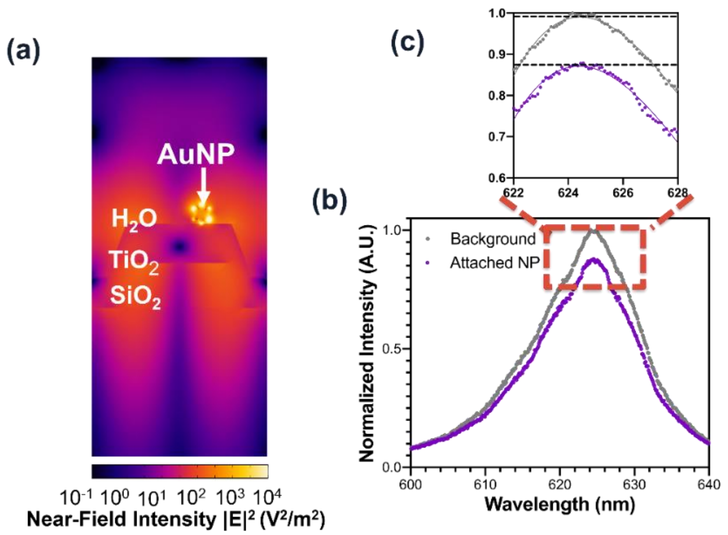


**Figure S2.** (a) Near-field profile of PC-AuNPs hybrid under resonant condition (λ = 620 nm at normal incidence). (b) Representative PC resonant reflected spectrum and (c) a close-up view of the reflected spectrum for the PC surface without or with attached AuNP. The reflected resonant intensity was obtained under a 20× lens.

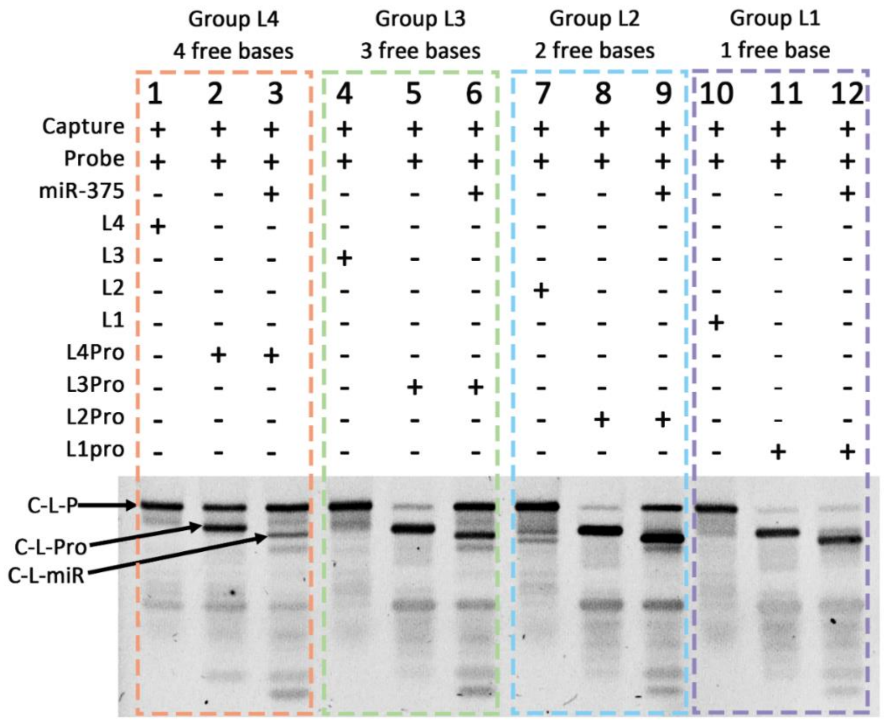


**Figure S3.** Native PAGE imaging of different length linker strands tested in the reaction.

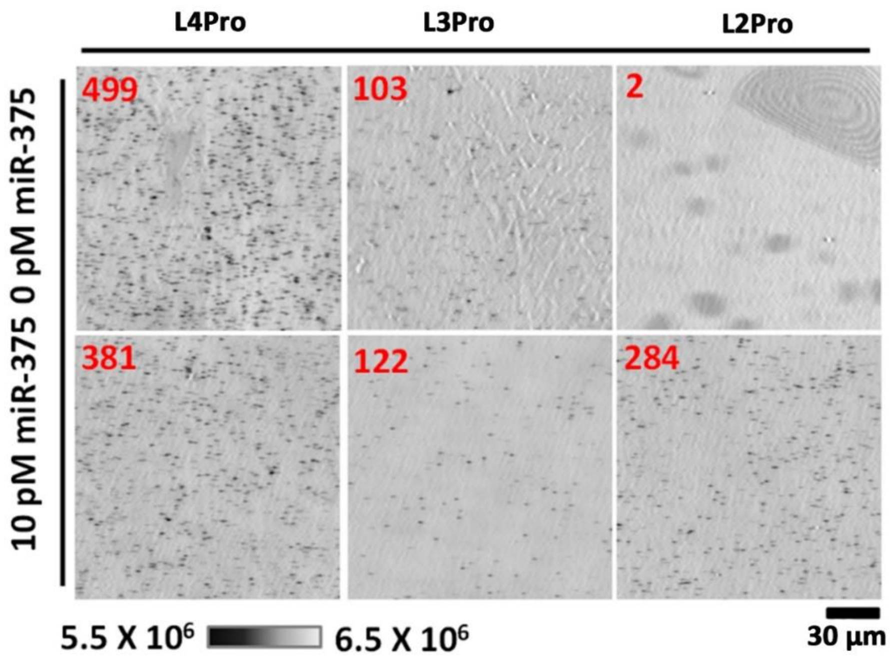


**Figure S4.** Optimization of linker strand length. TRAP image panel demonstrates particle count for different lengths of linker strand, captured at a reaction time of 30 minutes.

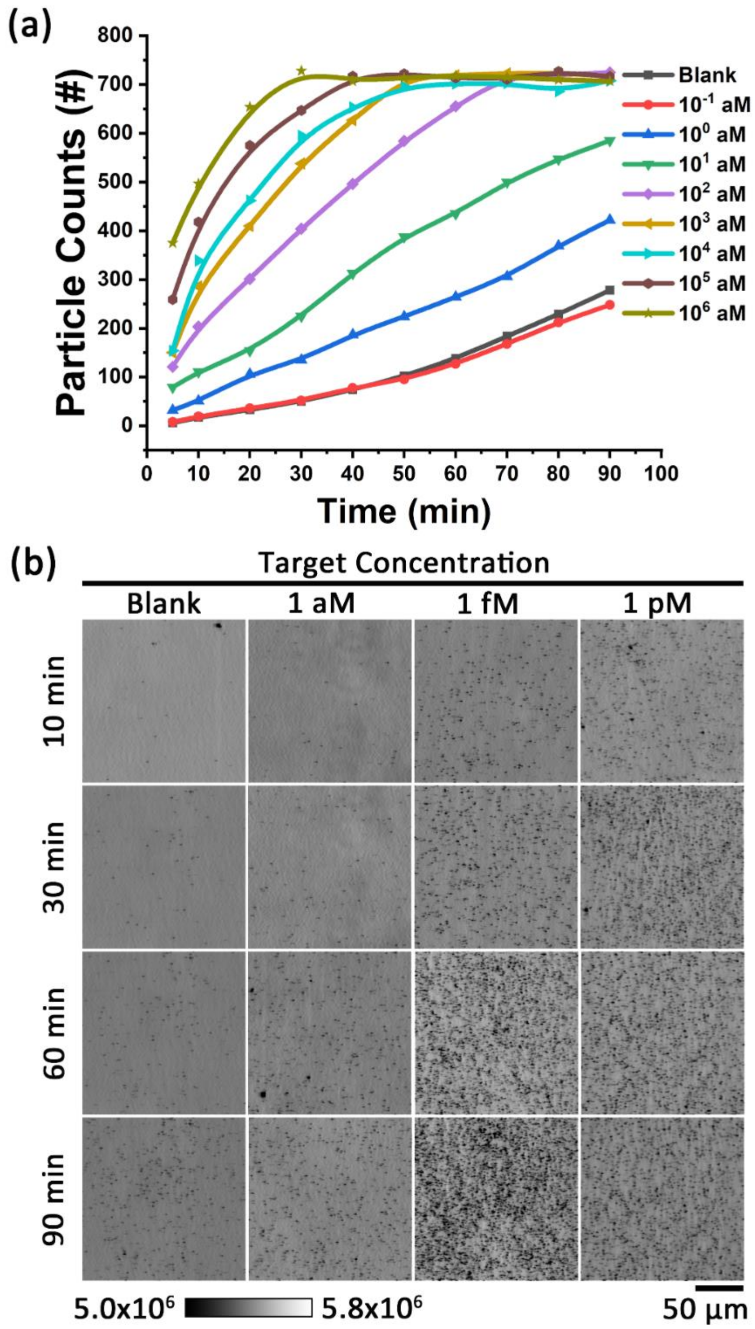


**Figure S5**. Kinetic discrimination of miR-375 concentration using PRAM and TRAP. (a) TRAP tests each concentration of the target miRNA miR-375 for analysis at 10 mins intervals over 90 mins. (b) Dose-response PRAM images of the TRAP assay at single particle resolution at 10 mins, 30 mins, 60 mins and 90 mins. Blank represents reaction without miRNA target present.

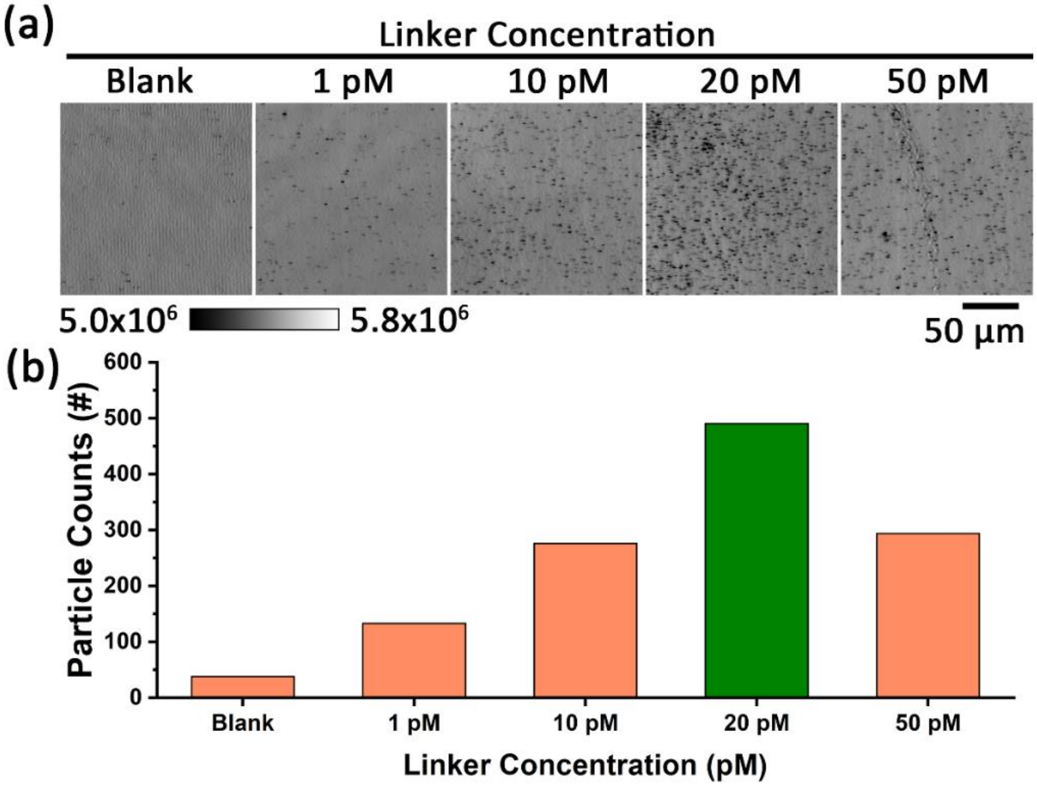


**Figure S6.** (a) Optimization of linker strand concentration imaged at 20 minutes. (b) particle count of various linker strand concentrations. Blank represents reaction without miRNA target present.

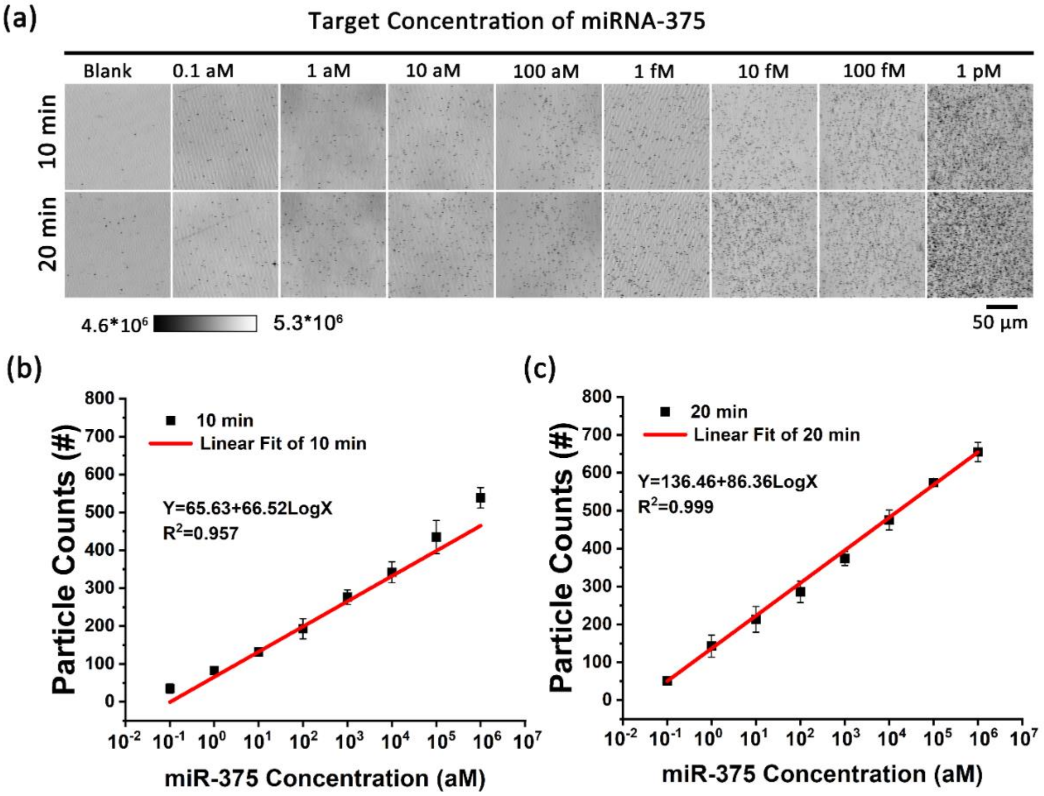


**Figure S7.** (a) TRAP image panel with digital resolution of captured AuNPs as a function of target concentration (rows) with 10 minutes and 20 minutes. Linear calibration curve of the miR-375 detection assay in buffer. (b) and (c) A linear regression model y = a + b × Log (x) was used to plot the dose response, where x is the concentration of target sequence, y is the AuNPs counts. (b) For the 10-minute response case, the linear regression equation was y=65.6+66.5 Log(x) with $R^2$ = 0.9573. (c) For the 20- minute response case, the linear regression equation was y=136.5+86.4 Log(x) with $R^2$ = 0.999. The dashed horizontal line indicates the threshold (blank signal + 3 standard derivations). The error bars represent the standard deviation of three independent assays.

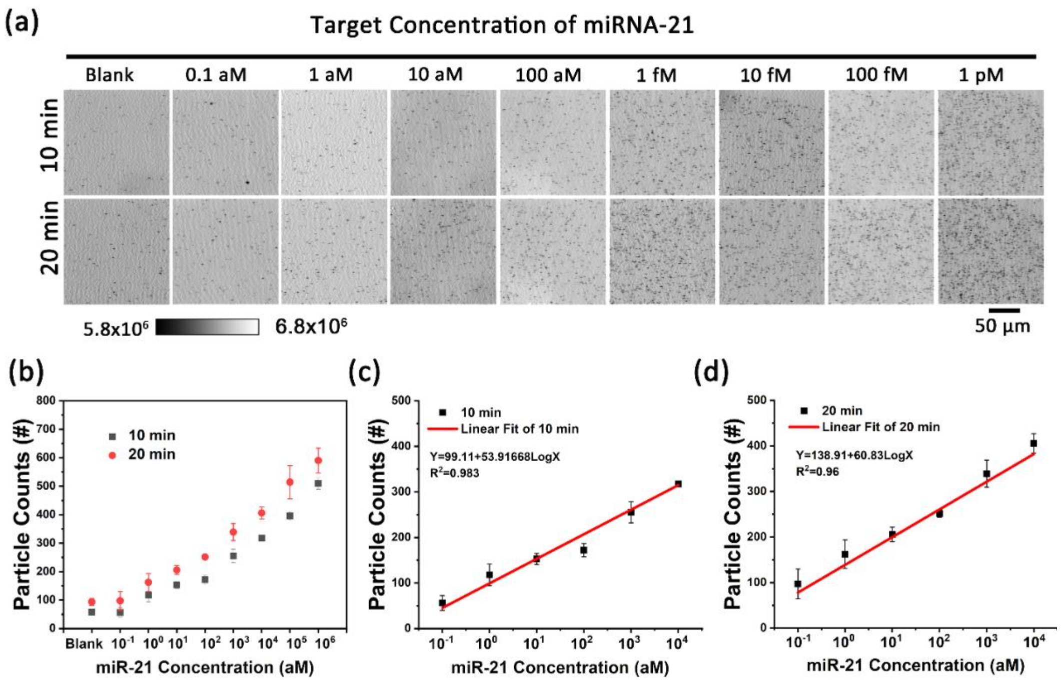


**Figure S8.** (a) TRAP image panel with digital resolution of captured AuNPs as a function of target concentration (rows) with 10 minutes and 20 minutes. Linear calibration curve of the miR-21 detection assay in buffer. (b) Quantification of particle count as a function of target concentration at 10 and 20 minutes. The blank represents the no-target control. (c) and (d) A linear regression model y = a + b × Log (x) was used to portrait the dose response, where x is the concentration of target sequence, y is the AuNPs counts. (b) For the 10-minute response case, the linear regression equation was y=96.54+43.52 Log(x) with $R^2$ = 0.982. (c) For the 20-minute response case, the linear regression equation was y=123.1+47.21 Log(x) with $R^2$ = 0.942. The dashed horizontal line indicates the threshold (blank signal + 3 standard derivations). The error bars represent the standard deviation of three independent assays.

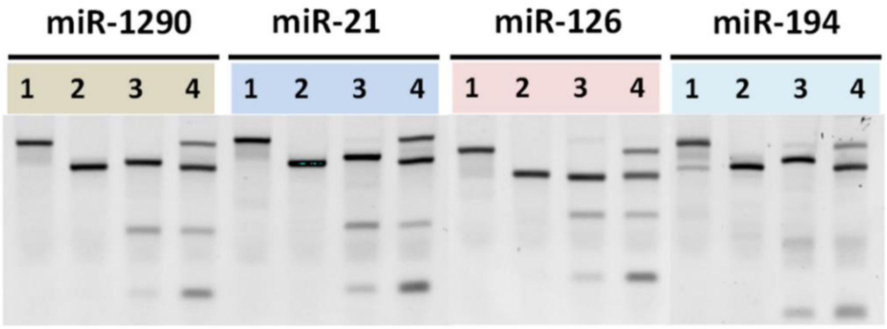


**Figure S9.** 12% PAGE. Lane 1: Capture + Linker + Probe; Lane 2: Capture + Linker +miRNA; Lane 3: Capture + Linker-Protector duplex + Probe; Lane 4: Capture + Linker-Protector duplex + Probe + miRNA;

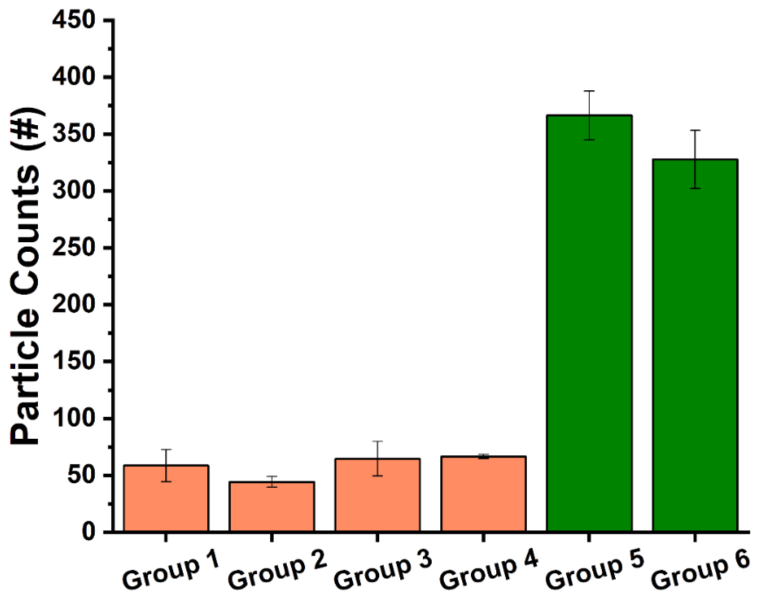


**Figure S10.** Responses of miRNA sensors to various controls at 15 mins. Group 1: miRNA-375 NPs and miRNA-375 LP with 1 pM target miRNA-21. Group 2: miRNA-375 NPs and miRNA-21 LP with 1 pM target miRNA-21. Group 3: miRNA-21 NPs and miRNA-21 LP with 1 pM target miRNA-375. Group 4: miRNA-21 NPs and miRNA-21 LP with 1 pM mixture target miRNA (miRNA-375, miRNA-126, miRNA 1290, miRNA-194). Group 5: miRNA-21 NPs and miRNA-21 LP with 1 fM miRNA-21 and 1 pM mixture target miRNA (miRNA-375, miRNA-126, miRNA 1290, miRNA-194). Group 6: miRNA-375 NPs and miRNA-375 LP with 1 fM miRNA-375 and 1 pM mixture target miRNA (miRNA-21, miRNA-126, miRNA 1290, miRNA-194).

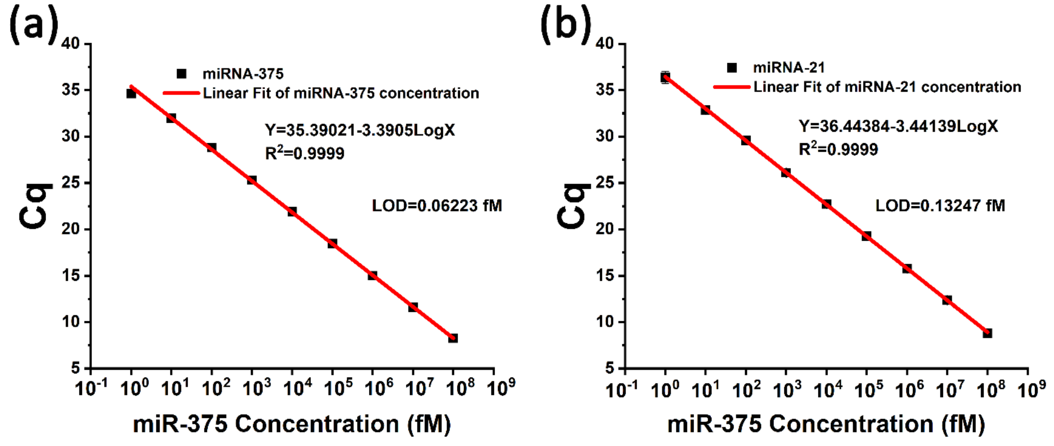


**Figure S11.** Linear calibration curve of the miR-375 and miR-21 detection assay in buffer by using qRT-PCR. (a) and (b) A linear regression model y = a + b × Log (x) was used to portrait the dose response, where x is the concentration of target sequence, y is the Cq. (b) For the miR-375 response case, the linear regression equation was y=35.39-3.39 Log(x) with $R^2$ = 0.9999. (c) For miR-21 response case, the linear regression equation was y=36.44-3.44 Log(x) with $R^2$ = 0.9999. The dashed horizontal line indicates the threshold (blank signal + 3 standard derivations). The error bars represent the standard deviation of three independent assays.

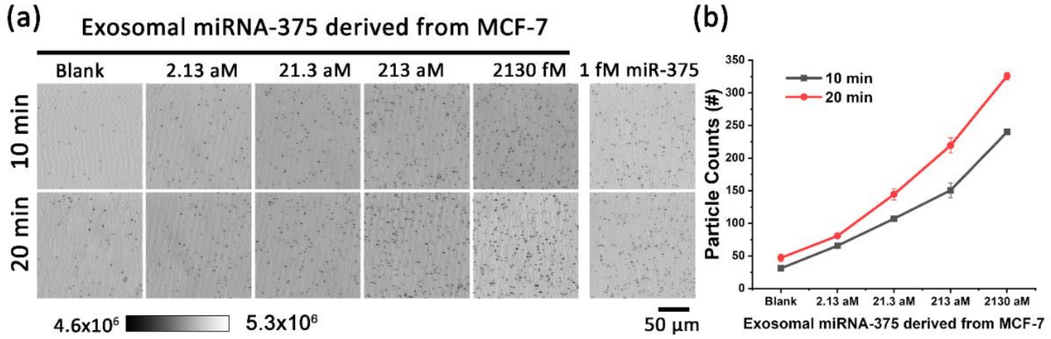


**Figure S12.** Kinetic discrimination of miR-375 concentration in MCF-7 cell exosomes using TRAP. (a) Dose-response plot for detection of miRNA-375 by TRAP, with detection shown at 10 and 20 minutes in a room temperature assay protocol. (b) Quantification of particle count as a function of miR-375 concentration in MCF-7 cell exosomes for three trials. The blank represents the no-target control. All PCs used for exosomal miRNAs were first validated with synthetic miRNA in buffer and normalized if necessary.

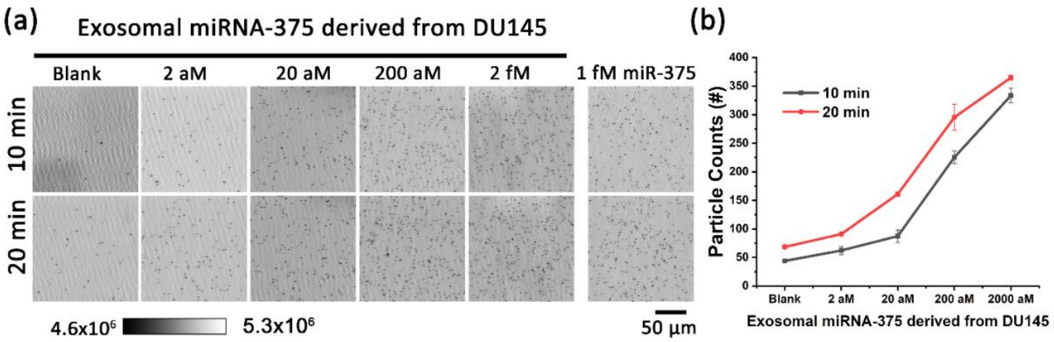


**Figure S13.** Kinetic discrimination of miR-375 concentration in DU145 cell exosomes using TRAP. (a) Dose-response plot for detection of miRNA-375 by TRAP, with detection shown at 10 and 20 minutes in a room temperature assay protocol. (b) Quantification of particle count as a function of miR-375 concentration in DU145 cell exosomes for three trials. The blank represents the no-target control. All PCs used for exosomal miRNAs were first validated with synthetic miRNA in buffer and normalized if necessary.

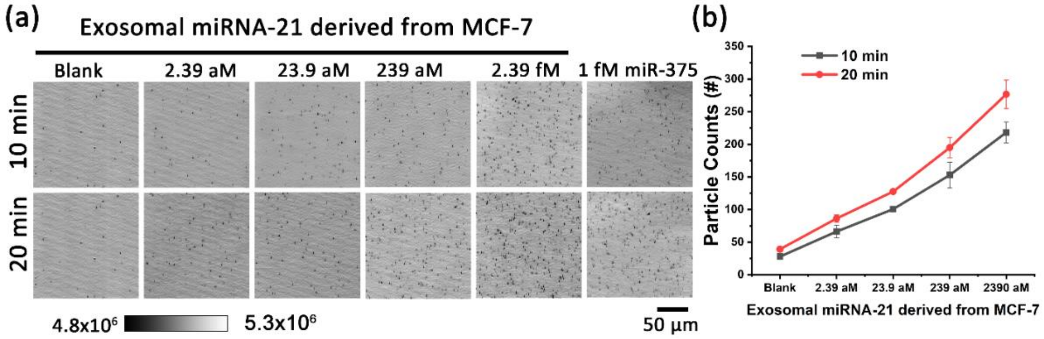


**Figure S14.** Kinetic discrimination of miR-21 concentration in MCF-7 cell exosomes using TRAP. (a) Dose-response plot for detection of miRNA-21 by TRAP, with detection shown at 10 and 20 minutes in a room temperature assay protocol. (b) Quantification of particle count as a function of miR-375 concentration in MCF-7 cell exosomes for three trials. The blank represents the no-target control. All PCs used for exosomal miRNAs were first validated with synthetic miRNA in buffer and normalized if necessary.

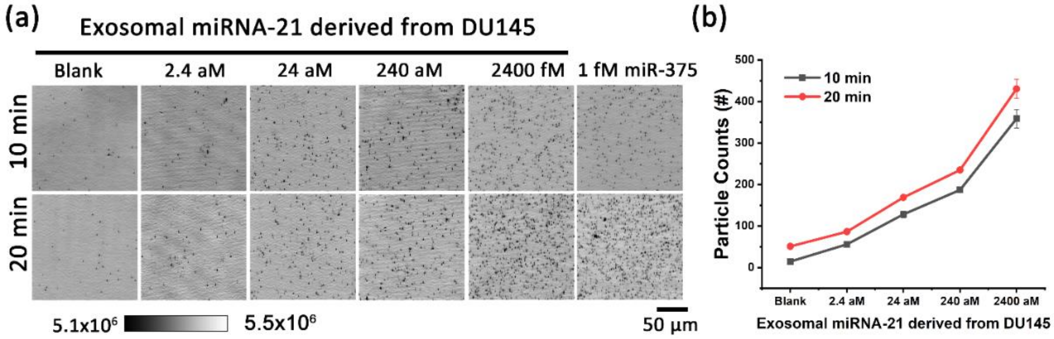


**Figure S15.** Kinetic discrimination of miR-21 concentration in DU145 cell exosomes using TRAP. (a) Dose-response plot for detection of miRNA-21 by TRAP, with detection shown at 10 and 20 minutes in a room temperature assay protocol. (b) Quantification of particle count as a function of miR-375 concentration in DU145 cell exosomes for three trials. The blank represents the no-target control. All PCs used for exosomal miRNAs were first validated with synthetic miRNA in buffer and normalized if necessary.

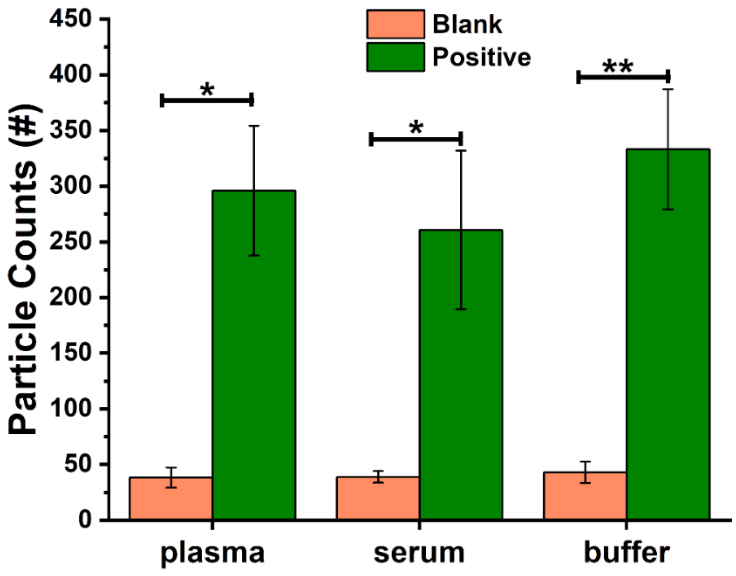


**Figure S16.** With miRNA-375 spiked in human plasma, serum, and buffer, TRAP counted the number of bound nanoparticles on the PC surface. Positive and blank represent reactions with 1 fM or absence of miRNA target, respectively. (Two-tailed Mann-Whitney t test)

[1] Y. Zhuo, H. Hu, W. Chen, M. Lu, L. Tian, H. Yu, K. D. Long, E. Chow, W. P. King, S. Singamaneni and B. T. Cunningham, *Analyst* **2014**, *139*, 1007-1015.

[2] D. Y. Zhang, S. X. Chen and P. Yin, *Nat. Chem.* **2012**, *4*, 208-214.
